# Gestational low-protein intake enhances the whole kidney miR-192 and miR-200 family expression and epithelial-to-mesenchymal transition in adult male offspring

**DOI:** 10.1101/199109

**Authors:** Letícia B Sene, Victor Hugo Gonçalves Rizzi, José AR Gontijo, Patricia A Boer

## Abstract

**Background:** Studies have been showed that maternal protein-restricted adult offspring, present pronounced reduction of nephron number associated with decreased fractional urinary sodium excretion and arterial hypertension. Also, recent advances in our understanding of the molecular pathways that govern the association of gestational nutritional restriction, intrauterine growth retardation inflammation with impaired nephrogenesis, nephron underdosing and kidney fibrosis point to the epithelial to mesenchymal transition (EMT) as the common.

**Method:** In the current study, the protein and sodium urinary excretion rates were evaluated and immunohistochemistry and western blot techniques were used to characterize the whole kidney structure changes in 16-wk old male LP offspring compared with age-matched controls. We also verify the expression of miRNAs, mRNAs and proteins markers of fibrosis and epithelial-to-mesenchymal transition in entire kidney prepared from LP offspring.

**Results:** In the current study, we may assume that arterial hypertension and long-term hyperfiltration process manifests, itself by proteinuria was accompanied by increased whole kidney mRNA expression of TGF-β1, ZEB1, type I collagen and, fibronectin in parallel to decreased expression of E-cadherin in 16-wk old LP offspring. Surprisingly, the renal tissue miR-129, miR-141, miR-200c and miR-429 were significantly upregulated in LP offspring compared to age-matched NP rats.

**Conclusion:** Considering that the overload in remaining nephrons, we may state that hypertension and proteinuria development following maternal protein restriction, may be a preponderant factor for the development of EMT and fibrous process and altered kidney ultrastructure in rat offspring. However, further studies are required to assess the contribution of miRNAs on renal injury progression in gestational protein-restricted model of fetal programming.

## INTRODUCTION

The gestational development is highly sensible to disturbances in the mother environment. Thus, an adverse intrauterine environment may alter the normal development, promoting adaptive growth restriction and lower birthweight. In addition, metabolic profile is programmed increasing the risk to disease development in adulthood [1,2]. The Developmental Origins of Health and Disease (DOHaD) concept define those gestational conditions that determinate changes to optimize the phenotype for adaptation in to postnatal environment. This process is known as fetal programming, with seminal impact to developmental biology [2,3].

Previous studies in our laboratory have demonstrated that gestational protein restriction is associated with renal morphological and physiological changes. Previous results from our laboratory showed that experimental protein restriction result in intrauterine growth retardation associated with impaired nephrogenesis and nephron underdosing [4-7]. These authors have correlated this abnormality with abnormal renal nerve activity, reduced nephron number and/or decreased glomerular filtration area in animals with a higher propensity to arterial hypertension, glomerulosclerosis and chronic renal failure (CKD). Thus, in this gestational programming animal model, we may suppose that the effective reduction of glomerular filtrated area promotes hyperflow and increased ultrafiltration pressure as a trigger to progressive sclerosis of glomeruli associated with tubular function disorders [4,5].

Epigenetic processes that include the control of gene expression by microRNAs orchestrate the intrauterine development. MicroRNAs (miRs) are small non-coding RNA that post-transcriptionally modulates gene expression by binding to the 3’ UTR of target mRNAs [8,9]. In this way, studies have demonstrated that miRs are involved in many biological processes including gene regulation, cell differentiation and proliferation, and, apoptosis [10,11].

The miR-200 family (including miR-200a, miR-200b, miR-200c, miR-141, and miR-429) and miR-192 are involved in epithelial-to-mesenchymal transition (EMT) biological process control [12-14]. EMT is an essential biological phenomenon that comprises a series of epithelial plasticity between the epithelial and mesenchymal states [15,16]. During EMT, the epithelial cells lose cell-cell contacts acquiring mesenchymal phenotype [17]. In addition, the phenotypic changes are associated with alterations in the transcription factors gene expression accompanied by decreased epithelial and enhanced expression of mesenchymal markers such as, respectively, E-cadherin and fibronectin [15]. EMT plays a fundamental role on pathophysiological processes of the renal fibrosis, a final common pathway that leads to CKD [16].

Whatever kidney pathological process, the renal interstitial fibrosis is common finding associated with severe kidney structure disorder and consequently, functional impairment [18]. Beta transforming growth factor (TGF-β) triggers the tubular EMT, and its expression is up-regulated in several type of chronic kidney disease [19]. Conversely, the miR-200f downregulates the kidney EMT expression induced by TGF-β1, via reduction of the translation of zinc finger E-box-binding homeobox1/2 transcription factors (ZEB1 and ZEB2).

Recently, in the isolated glomeruli study, we found that male adult offspring from gestational protein-restricted mothers, present reduced miR-200f expression associated with enhanced TGF-β1 and ZEB2 expression, which are related to glomeruli EMT and glomerular fibrosis markers [20]. The current study was performed to evaluate the miR-192 and miR-200f expression as well as the EMT occurrence, in whole kidney tissue from maternal protein-restricted adult offspring. Surprisingly, here we found a distinct pattern of miRNAs expression in whole kidney tissue when compared to isolated glomeruli from these programmed animals.

## MATERIALS AND METHODS

### Animals and experimental design

The experiments were conducted on female and male age-matched *Wistar HanUnib* sibling-mated rats (250–300g) obtained from colonies maintained under specific pathogen-free conditions in the Multidisciplinary Center for Biological Investigation CEMIB/Unicamp, Campinas, Brazil. The environment and housing presented good conditions for managing their health and well-being during the experimental procedure. The study design was approved by the São Paulo State University Institutional Animal Ethics Committee (protocol CEUA/UNESP #292) and conformed to general guidelines established by the Brazilian College of Animal Experimentation (COBEA). Immediately after weaning at 3 weeks of age, animals were maintained under controlled temperature (25°C) and lighting conditions (0700h-1900h), with free access to tap water and standard rodent laboratory chow. We designated as day 1 of pregnancy the day in which the vaginal smear presented sperm. The dams were maintained on isocaloric rodent laboratory chow with normal protein content [NP], (17% protein) or low protein content [LP] (6% protein) diet, *ad libitum* intake, throughout the entire pregnancy. All dams groups returned to the NP diet intake immediately after delivery. The offspring birthweight was measured. The pups weaned in 3 weeks and only one male offspring of each litter was used for each experiment. The male offspring were maintained on standard rodent laboratory with normal protein content under a controlled room temperature and lighting conditions and, followed up to 16 weeks of age. Food consumption was monitored daily and normalized to the body weight. The systolic arterial pressure was measured in conscious 16-week-old offspring (NP n=9 and LP n=8) by an indirect tail-cuff method using an electrosphygmomanometer (IITC Life Science – BpMonWin Monitor Version 1.33) combined with a pneumatic pulse transducer/amplifier. This indirect approach allowed repeated measurements with a close correlation (correlation coefficient = 0.975) compared to direct intra-arterial recording. The mean of three consecutive readings represented the blood pressure. Body weight was recorded weekly. Twelve-day-old and sixteen week-old male offspring were euthanized and the kidneys collected for real-time PCR, western blot, Sirius red and immunohistochemistry analyzes.

### Renal function test and measurement of Proteinuria

The renal function tests were performed on the last day at 16 weeks of age in unanaesthetized, unrestrained NP (n = 10) and LP (n = 11) male offspring. Creatinine clearance was done as standard methodology [4-6]. Briefly, after an overnight fast, each animal received a load of tap water by gavage (5% of body weight), followed by a second load of the same volume, 1 hour later, and spontaneously voided urine was collected over a 120-min period into a graduated centrifuge tube Spontaneously voided urine was collected over a 120-min period into a centrifuge tube and measured gravimetrically. At the end of the experiment, blood samples were drawn through cardiac puncture in anesthetized rats, and urine and plasma samples were collected for analysis [3-6,31,32]. Plasma and urine sodium and potassium concentrations were measured by flame photometry (Micronal, B262, São Paulo, Brazil), while the creatinine concentrations was determined spectrophotometrically (Instruments Laboratory, Genesys V, USA). The proteinuria was detected using the Sensiprot Kit (Labtest).

### Total RNA extraction

RNA was extracted from entire kidney (n=5 for each group from 5 different mothers) using Trizol reagent (Invitrogen), according to the instructions specified by manufacturer. Total RNA quantity was determined by the absorbance at 260 nm using nanoVue spectrophotometer (GE healthcare, USA), and the RNA purity was assessed by the A 260nm/A 280nm and A 260nm/A 230nm ratios (acceptable when both ratios were > 1.8). RNA Integrity was ensured by obtaining a RNA Integrity Number - RIN>8 with Agilent 2100 Bioanalyzer (Agilent Technologies, Germany).

### Reverse Transcription of miRNA and mRNA

cDNA was synthesized using TaqMan^®^microRNA Reverse Transcription kit (Life Technologies, USA), combined with Stem-loop RT Primers (Life Technologies, USA) and High Capacity RNA-to-cDNA Master Mix (Life Technologies, USA) according the manufacturer’s guidelines. For miRNA, 3µl (10ng) total RNA was mixed with specific primers (3µl), dNTPs (100mM), MultiScribe TM Reverse Transcriptase (50µl), 10X RT Buffer, RNase inhibitor (20µl) and completed up to 4.5µl with H_2_O. The cycling conditions were 16°C for 2-min, 42°C for 1min, 50 °C for 1-second and 85°C for 5-min. For mRNA, 10-µl total RNA was mixed with 4µl Master Mix, 2µl specific primers and completed up to 20µl with H_2_O. The cycling conditions were 25°C for 5 min, 42°C for 30 min and 85°C for 5-min.

### Real-time quantitative PCR (miRNAs)

Each cDNA of miRNA-200 family (miR-200a, miR-200b, miR-200c, miR-141 and miR-429) and miR-192 was quantified by real-time quantitative PCR using ABI Prism 7900 Sequence Detection System (Life Technologies, USA). We used, for each reaction, 10µl TaqMan ^®^ Universal PCR Master Mix, 2µl TaqMan MicroRNA Assay Mix (Life Technologies, USA), and 1.5µl cDNA and completed up to 20µl reaction volume. The cycling conditions were 95°C for 10 minutes; 45 cycles of 95°C for 15 seconds and 60°C for 1-minute.

### Real-time quantitative PCR (mRNAs)

For the analysis of expression level of ZEB1, ZEB2, desmin, fibronectin, ZO-1, E-cadherin, TGF-β1, col 1α1 and col 1α2, in renal tissue, RTqPCR was carried out with SYBR green Master Mix, using specific primers for each gene (Table 1). Reactions were set up in a total volume of 20µL using 5µl of cDNA (diluted 1:100), 10µL SYBR green Master Mix (Life Technologies, USA) and 2.5µL of each specific primer (5nM) and performed in the ABI Prism 7300 real-time PCR system (Life Technologies, USA). The cycling conditions were 95°C for 10-minutes; 45 cycles of 95°C for 15-sec and 60°C for 1-min.

**Table 1.**
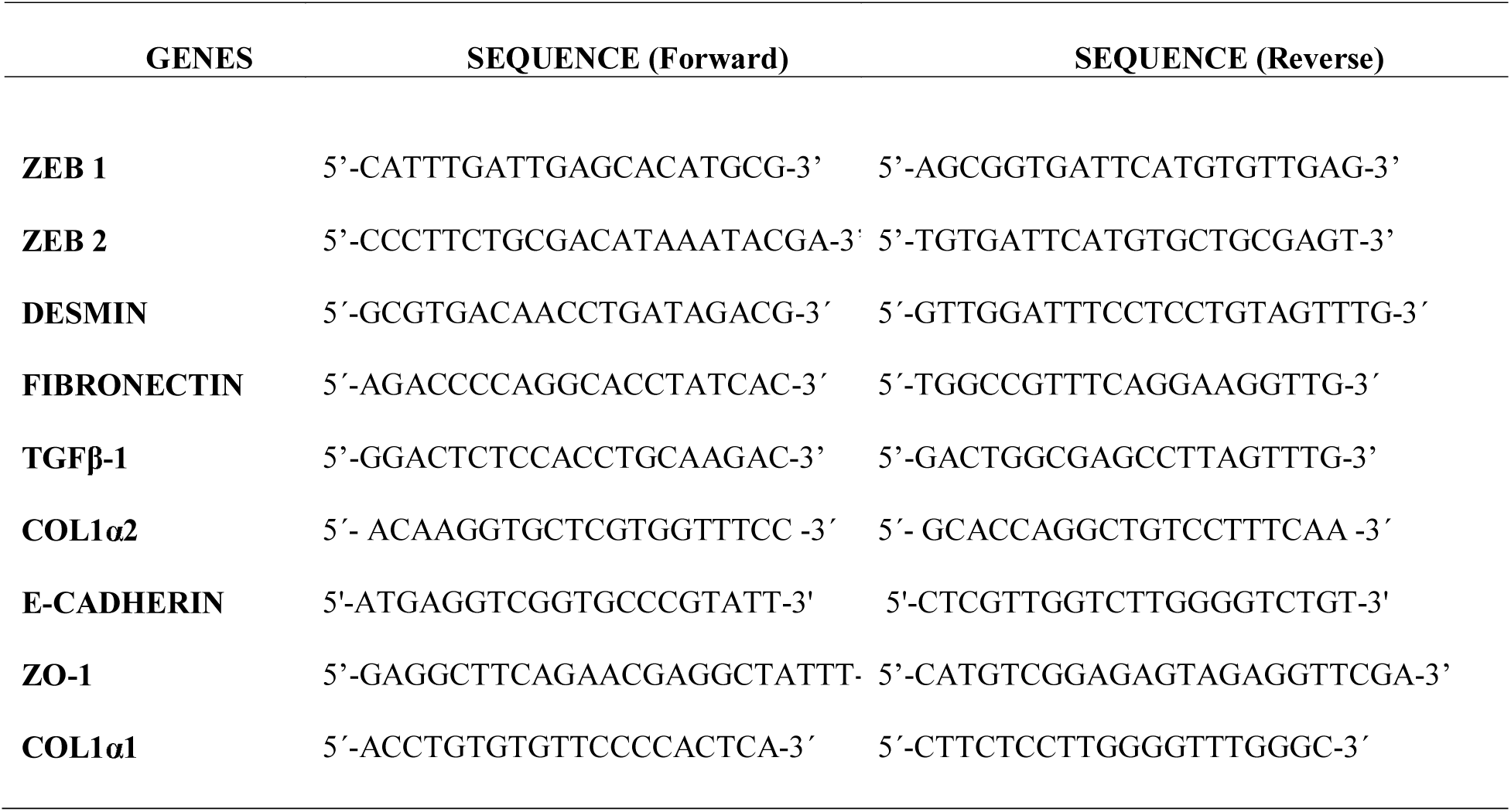
Gene sequence studied in whole kidney from 12 days and 16 week-old in LP compared to age matched NP offspring.

### Analysis of the gene expression

To analyze the differential expressions, the miRNA or mRNA levels obtained for each gene (Table 1) were compared with LP group with respect to the appropriated NP group. Normalization of miRNA expression was made using the expression of the snRNA U6 and snRNA U87 reference genes (Accession: NR_004394 and AF272707, respectively), and, for mRNA expression, the genes GAPDH, β-actin and TBP. Relative gene expression was evaluated using the comparative quantification method (PFAFFL, 2001). All relative quantifications were assessed using DataAssist software v 3.0, using the ∆∆CT method. PCR efficiencies calculated by linear regression from fluorescence increase in the exponential phase in the program LinRegPCR v 11.1 [21].

### Immunoblotting

The entire kidney tissue (n=5 for each group from 5 different mothers) was removed and frozen in liquid nitrogen and stored at -80°C. After, the tissue was minced coarsely and homogenized immediately in 10 volumes of solubilization buffer (10 ml/L Triton-X 100, 100mM Tris[hydroxymethyl]amino-methane (Tris) pH 7.4, 10mM sodium pyrophosphate, 100mM sodium fluoride, 10mM ethylendiaminetetracetic acid (EDTA), 10mM sodium vanadate, 2mM phenylmethylsulfonyl fluoride (PSMF) and 0.1mg/ml aprotinin at 4°C), using a polytron PTA 20S generator (model PT 10/35, Brinkmann Instruments, Westbury, N.Y., USA) operated at maximum speed for 20-sec. The tissue extracts were centrifuged at 11.000 rpm at 4°C for 40-min, and the supernatants were used as sample. Protein quantification was performed using the Bradford method. The samples were dissolved in Laemmli buffer, heated in a boiling water bath for 5-min., and were used for SDS-PAGE (Bio-Rad Laboratories, Hercules, CA, USA). Equal amounts of protein (100µg) of each sample were loaded per well onto preformed gradient gels 4-15% (Bio-Rad Laboratories) for 1hr 30min to 100v. After electrophoretic separation, proteins were transferred to nitrocellulose membranes and then blocked with TBS-T solution containing 5% non-fat dry milk at room temperature for 1-h. Nitrocellulose blots were then incubated at 4°C overnight with primary antibodies diluted in TBS-T or BSA 1% as follow: E-cadherin (ab53033, 1:200, Abcam), desmin (ab15200, 1:200, Abcam), ZEB1 (sc-10572, 1:200, Santa Cruz), ZEB2 (sc-48789, 1:200, Santa Cruz), TGFβ1 (sc-146, 1:100, Santa Cruz), and Collagen 1 (sc-8788, 1:100, Santa Cruz). Immunoreactivity bands were detected using the enhanced chemiluminescence substrate kit (Pierce ECL Western Blotting Substrate – GE Healthcare, Pittsburgh, PA, USA) and the images were obtained by CCD camera (G: BOX Chemi, Syngen^®^, Sacramento, CA, USA), and band intensities were quantified by optical densitometry (UN-SCAn-IT gel, Gel & Graph Digitizing Software version 6.1). Beta-actin was used as an endogenous control.

### Histology and immunohistochemistry (IHC)

Animals were deeply anaesthetized with a mixture of ketamine (75 mg/kg body weight, i.p.) and xylasine (10mg/kg body weight, i.p.) and the level of anesthesia was controlled by monitoring the corneal reflex. The rats were transcardially perfused with saline followed by paraformaldehyde 4% in 0.1 M phosphate buffer, pH 7.4. After perfusion, the kidneys (n =5 for each group from 5 different mothers) were removed, weighed and placed in the same fixative for 2h, followed by 70% alcohol until processed for paraffin inclusion. The paraffin blocks were cut into 5-µm-thickness sections. The picrosirius technique was used to evaluate the density of collagen. Ten cortical fields of histological sections (n = 5 for each group) were analyzed, and the average of the readings determined the density of collagen. Images were captured with a photomicroscope and analyzed by Leica Qwin 3.1 for Windows. For IHC, paraffin sections were incubated with primary antibodies, overnight at 4°C, for anti-collagen 1 (1:1000, Sigma), anti-fibronectin (1:000 Novo Castra), anti-TGF-β1 (1:000 Santa Cruz), anti-E-cadherin (1:000 Abcam), and anti-ZEB1 e 2 (1:000 Santa Cruz). Secondary antibodies were used according to the primary antibody. Finally, sections were revealed with 3, 3’- diaminobenzidine tetrahydrochloride (DAB – Sigma - Aldrich CO^®^, USA), counterstained with Mayer’s hematoxylin, dehydrated and mounted.

### Statistical Analysis

Data obtained from this study are expressed as the mean ± SD. Comparisons involving only two means within or between groups were carried out using a Student’s t-test. Statistical analysis was performed with GraphPad Prism 5.01 for Windows (1992-2007 GraphPad Software, Inc., La Jolla, CA, USA). The level of significance was set at *P* < 0.05.

## RESULTS

### Kidney and body weight

Gestational protein restriction did not significantly change the pregnant dams’ body mass during gestation. In addition, it did not affect the number of offspring per litter and the proportion of male and female offspring (p=0.3245). The birthweight of LP male pups (n=35) was significantly reduced compared with NP pups (n=32)(5.0 ± 0.09 g vs. 6.0 ± 0.06 g; p=0.0001) (Table 2). The body mass of 12 days weaning LP pups (21 ± 0.5 g, n=18) remained lower than compared age-matched NP pups (24 ± 0.6 g, n=19) (p=0.0017) (Table 2). However, at 16 weeks of age, the body weight of LP and NP rats was similar (p=0.3927). Additionally, at 12 days after birth, the both, left and right kidney weight of LP offspring was significantly reduced when compared to age-matched NP offspring; however, at 16-wk of life the kidney weight were similar in both, NP and LP offspring groups (Table 2).

**Table 2.** Offspring body and kidney weight (in grams) at birth, from 12 days and 16 week-old in LP compared to age matched NP offspring. The results are expressed as means ± SD. Student’s t-test

| **Body/Kidney weight** | **NP** | **LP** | ***p*** |
| --- | --- | --- | --- |
| ***At Birth*** |  |  |  |
| Birthweight | 6 ± 0.06 (n = 32) | 5 ± 0.09 (n = 35) | 0.0001 |
| ***12-day old*** |  |  |  |
| Body weight (b.w.) | 24±0.6 (n=19) | 21±0.5 (n=18) | 0.0017 |
| Left kidney/b.w. | 0.007±0.0002 (n=12) | 0.006±0.0001 (n=11) | 0.009 |
| Right kidney/b.w. | 0.007±0.0002 (n=12) | 0.006±0.0001 (n=11) | 0.01 |
| ***16-wk old*** |  |  |  |
| Body weight (b.w.) | 456 ± 15 (n = 7) | 472 ± 10 (n = 7) | 0.3927 |
| Left kidney/b.w. | 0.003 ± 0.00008 (n = 6) | 0.003 ± 0.0001 (n = 6) | 0.21 |
| Right kidney/b.w. | 0.005 ± 0.0002 (n = 7) | 0.004 ± 0.0001 (n = 7) | 0.057 |

### Renal function data

The data for renal function in the 16-week-old offspring of both (NP and LP) groups are summarized in Table 3. There were no significant differences between serum sodium, potassium, lithium and creatinine levels in NP rats, compared with the LP group. The urinary flow rates (data not included) and the glomerular filtration rate, estimated by CCr, did not significantly differ among the groups during the renal tubule sodium handling studies (Table 3). Fractional urinary sodium excretion (FENa) was significantly lower in low maternal protein intake rats when compared with the normal maternal protein intake age matched group, as follows: LP: 0.96 ± 0.036% versus NP: 1.44 ± 0.26% (p ≤0.05). Urine from LP rats (45.92±19.6 mg/day, n□=□10, p=0.03) showed an elevated protein level when compared with age-matched NP rats (17.41±4.9 mg/day, n□=□10), as shown in Table 3.

**Table 3.** Arterial blood pressure in the 16-week-old LP compared to age-matched NP offspring (n=10 for each group). Creatinine clearance (*CCr*), Fractional urinary sodium excretion (*FENa+*) and proteinuria in the 16-wk-old LP compared to age-matched NP offspring (n=11 for each group).

| ***Groups*** | ***Blood Pressure***<br>***(mmHg)*** | ***CCr***<br>***ml/min/100g b.w.)*** | ***FENa+***<br>***(%)*** | ***Proteinuria***<br>***(mg/day)*** |
| --- | --- | --- | --- | --- |
| ***NP*** | 126.5±8.4 | 1.20±0.2 | 1.44±0.26 | 17.41±4.9 |
| ***LP*** | 139.3±5.7* | 1.04±0.2 | 0.96±0.036* | 45.92±19.6* |

The data are reported as the means ± SD. \* $P \leq 0.05$ versus LP (Student's t test).

### Expression profile of miR-192 and miR-200 family

In 12-day old offspring the renal LP expression of all studied miRs was not altered comparatively to that found in NP (Figure 1A). On the other hand, in 16-wk old LP animals, the renal tissue miR-129, miR-141, miR-200c and miR-429 were significantly upregulated when compared to that observed in age-matched NP. The expression of miR 200a/b was unchanged (Figure 1B).

**Figure 1.**
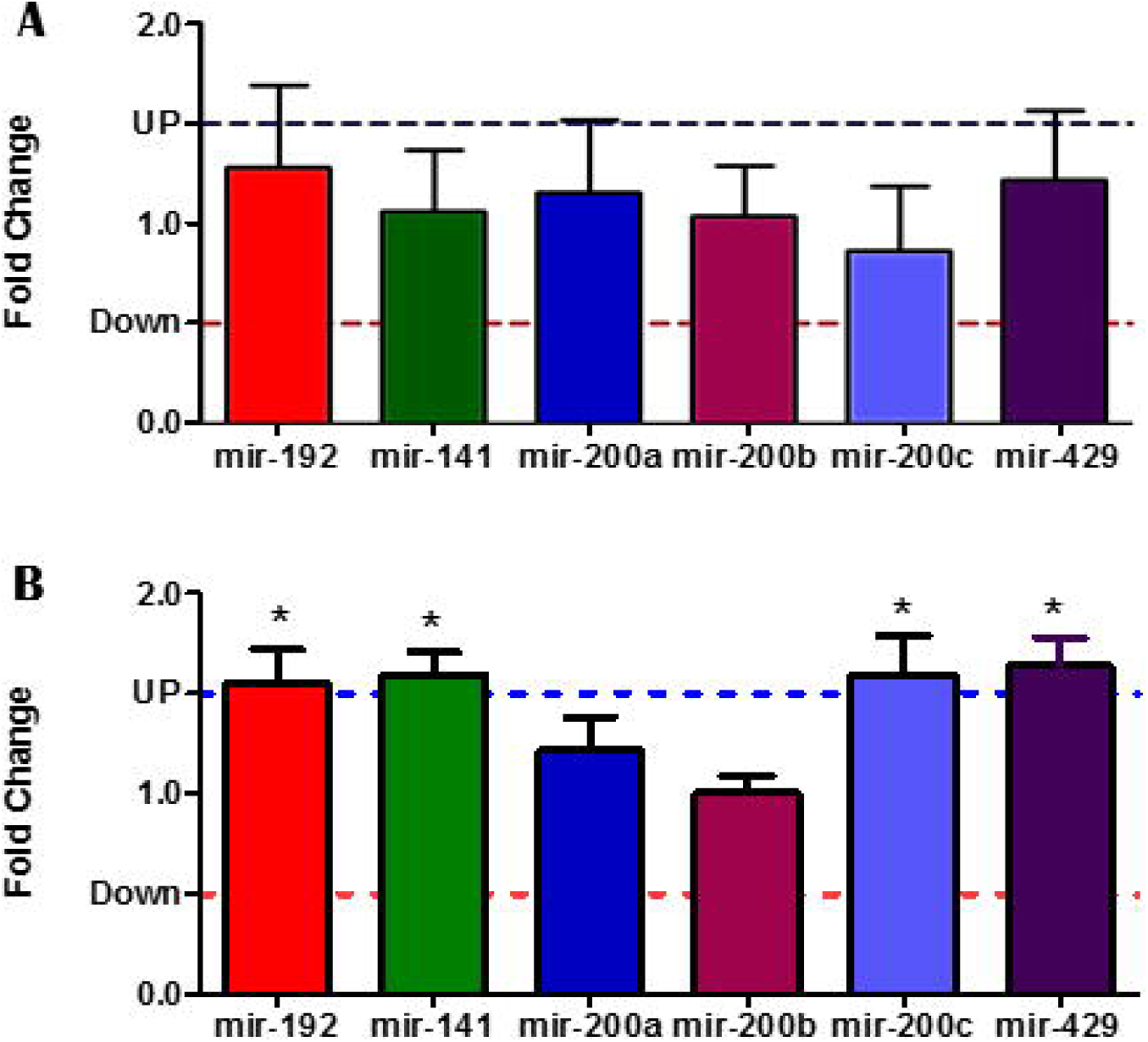
Expression of the miR-200 family and miR-192 from 12 days (A) and 16 week-old(B) in LP compared to NP offspring. The normalization of miRNA expression was made using the expression of the snRNA U6 and snRNA U87 reference genes (Accession: NR_004394 and AF272707, respectively). Relative gene expression was evaluated using the comparative quantification method.

### Gene expression

The gene (mRNA) expression to collagen 1α1, collagen 1α2 and ZEB1 was significantly increased in the whole kidneys from 16-wk-old LP, comparatively to age-matched NP offspring (Figure 2). At this age, desmin, E-cadherin, fibronectin, TGF-β, ZEB2 and ZO-1 mRNAs expression were not altered (Figure 2).

**Figure 2.**
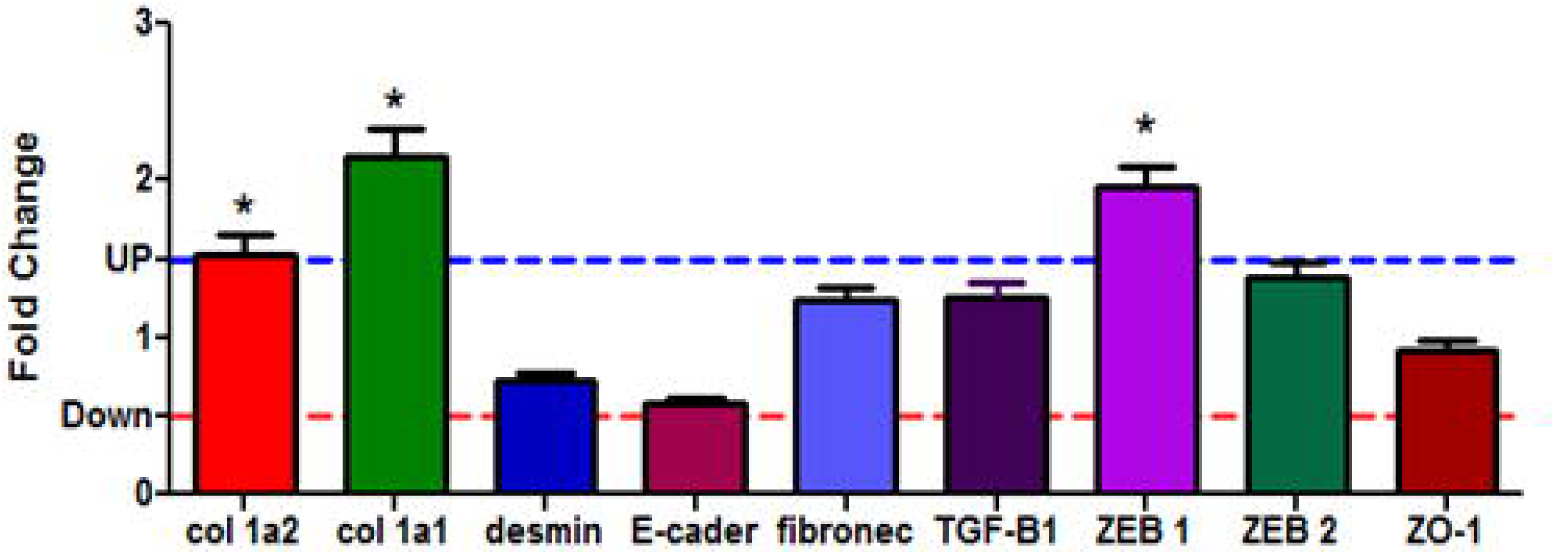
The differential gene (mRNA) expression to collagen 1α1, collagen 1α2, ZEB1, ZEB2, E-cadherin, fibronectin, TGF-β1, desmin and ZO-1 from kidneys in 16-wk-old LP, comparatively to age-matched NP offspring.

### Proteins analysis

The study of fibrosis markers and EMT-related proteins by both, immunohistochemistry and western blot analysis, shows a significant rise of the renal TGF-β1 expression in 16-wk-old LP comparatively to NP age-matched offspring (Figure 3). This was observed in both, renal cortical and medullar tissues. The expression of extracellular matrix proteins compounds was also enhanced in kidneys of 16-wk-old LP offspring. Using picrosirius estimation analysis, the current study shows a significant enhancement of collagen content in the kidneys of 16-wk-old LP offspring when compared with age-matched NP group (Figure 4A-C). Type I collagen immunoreactivity was enhanced in adult LP however, by western blot analysis, type I collagen expression was not statistically significant when compared both offspring groups (NP and LP) (Figure 4D-H). In addition, fibronectin immunoreactivity was enhanced in the renal cortical and medullar areas of 16-wk-old LP offspring (Figure 5). The entire kidney immunoreactivity to E-cadherin was decreased (Figure 6A and B) while ZEB1 was enhanced (Figure 6D and E) in adult LP when compared with age-matched NP offspring. The western blot semi-quantitative analyze confirm that significant difference showed in the immunohistochemical studies (figure 6C and F). We did not found any difference to desmin and ZEB2 expressions in LP compared to appropriate age-matched controls (Figure 7).

**Figure 3.**
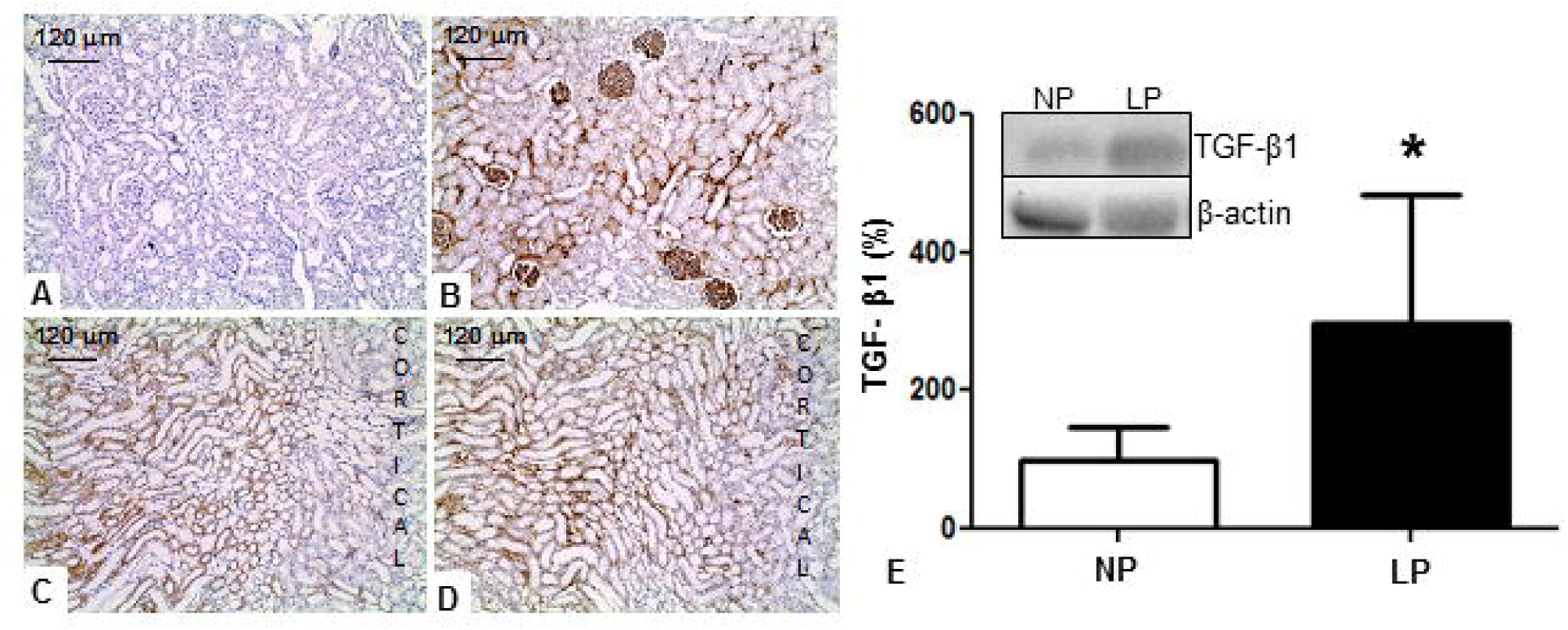
TGF-β1 immunohistochemistry and western blot expression of renal tissue from 16-wk-old LP (B, D and E), comparatively to age-matched NP (A, C and E) offspring.

**Figure 4.**
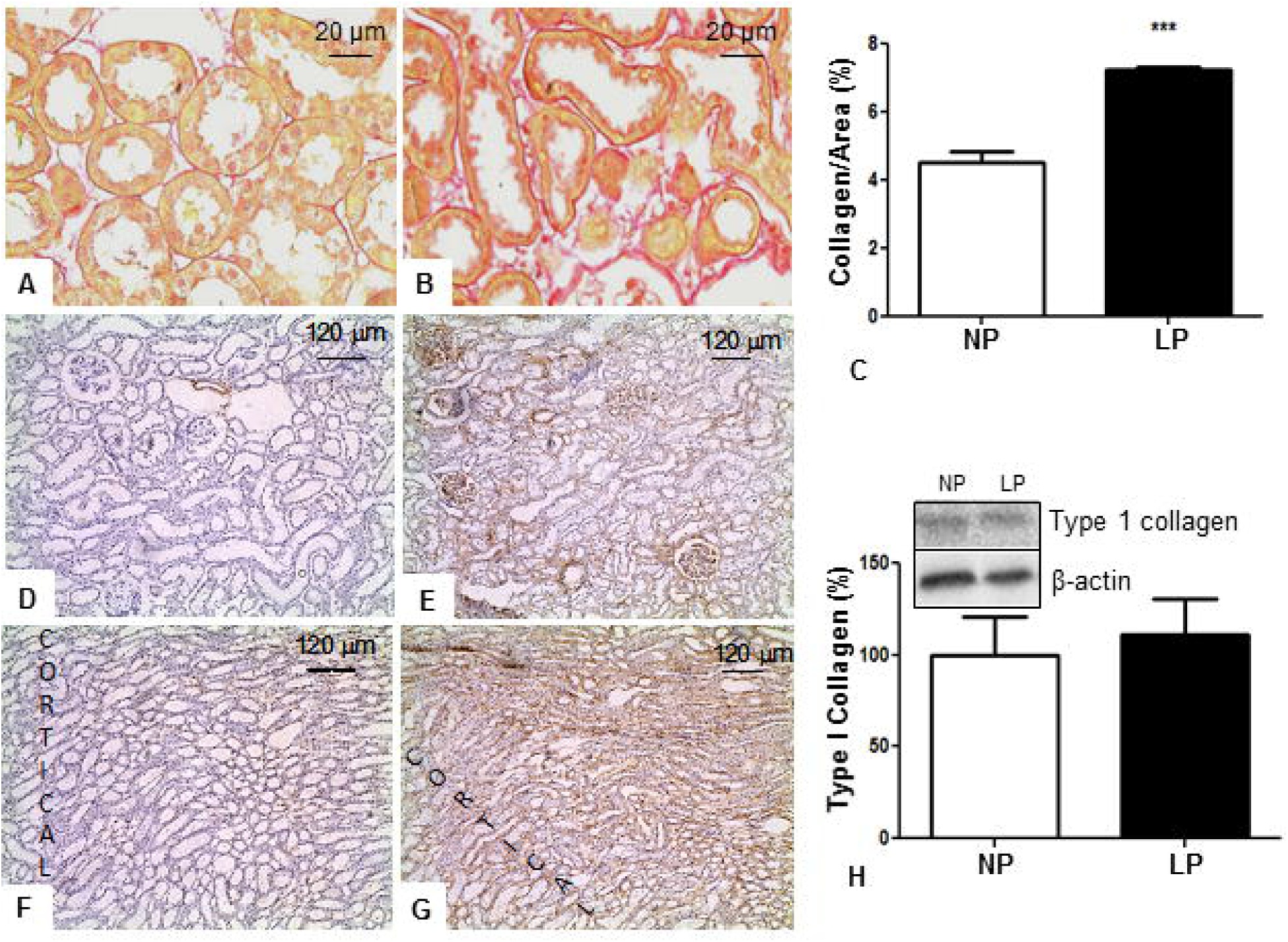
Quantification of picrosirius labeled area (C) in NP (A) and LP (B) renal tissue from 16-wk-old offspring. Type 1 collagen immunohistochemistry in NP (D and F) and LP (E andG) kidneys and western blot quantification (H).

**Figure 5.**
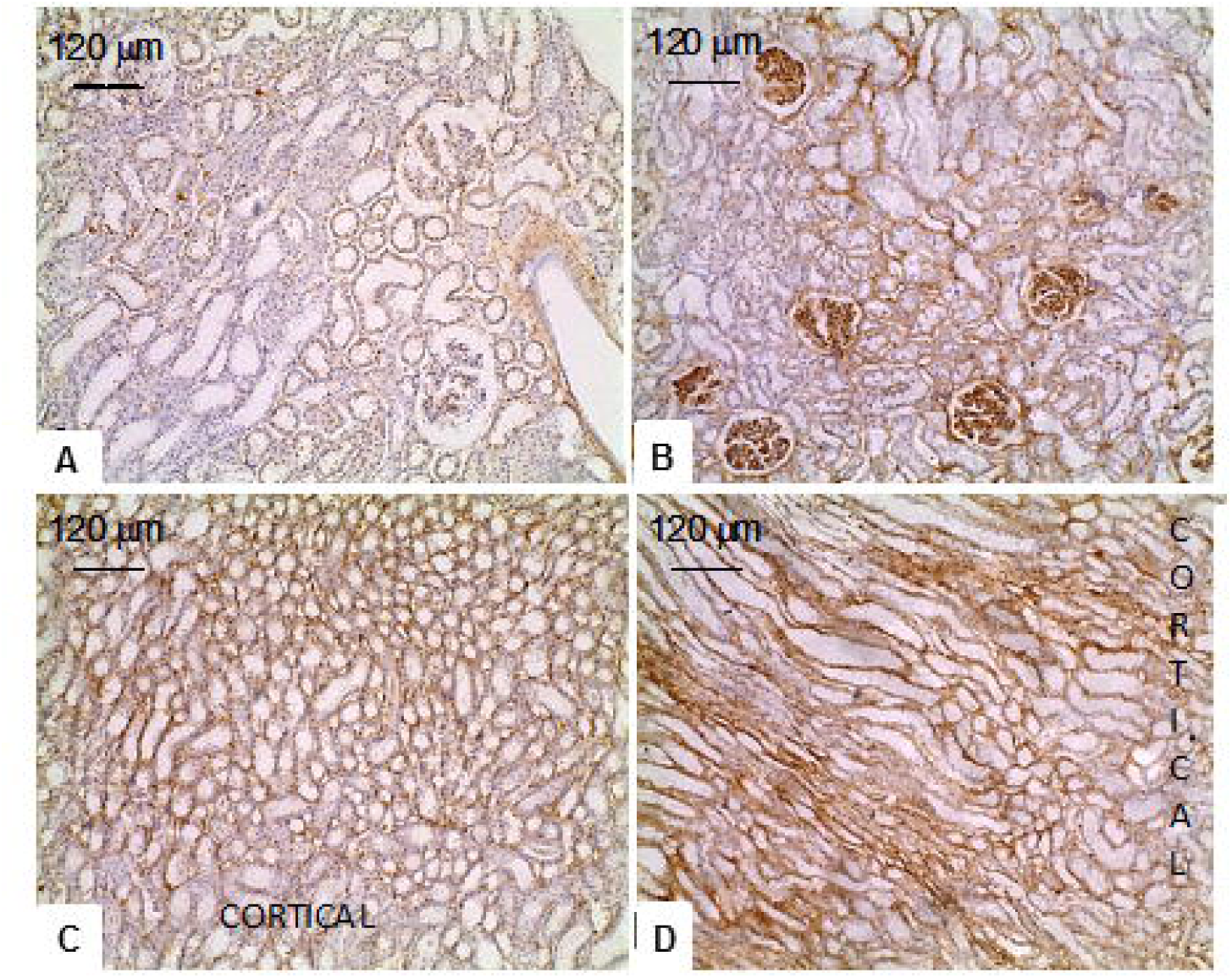
Fibronectin immunohistochemistry from 16-wk-old LP (B and D), comparatively to age-matched NP (A and C) offspring.

**Figure 6.**
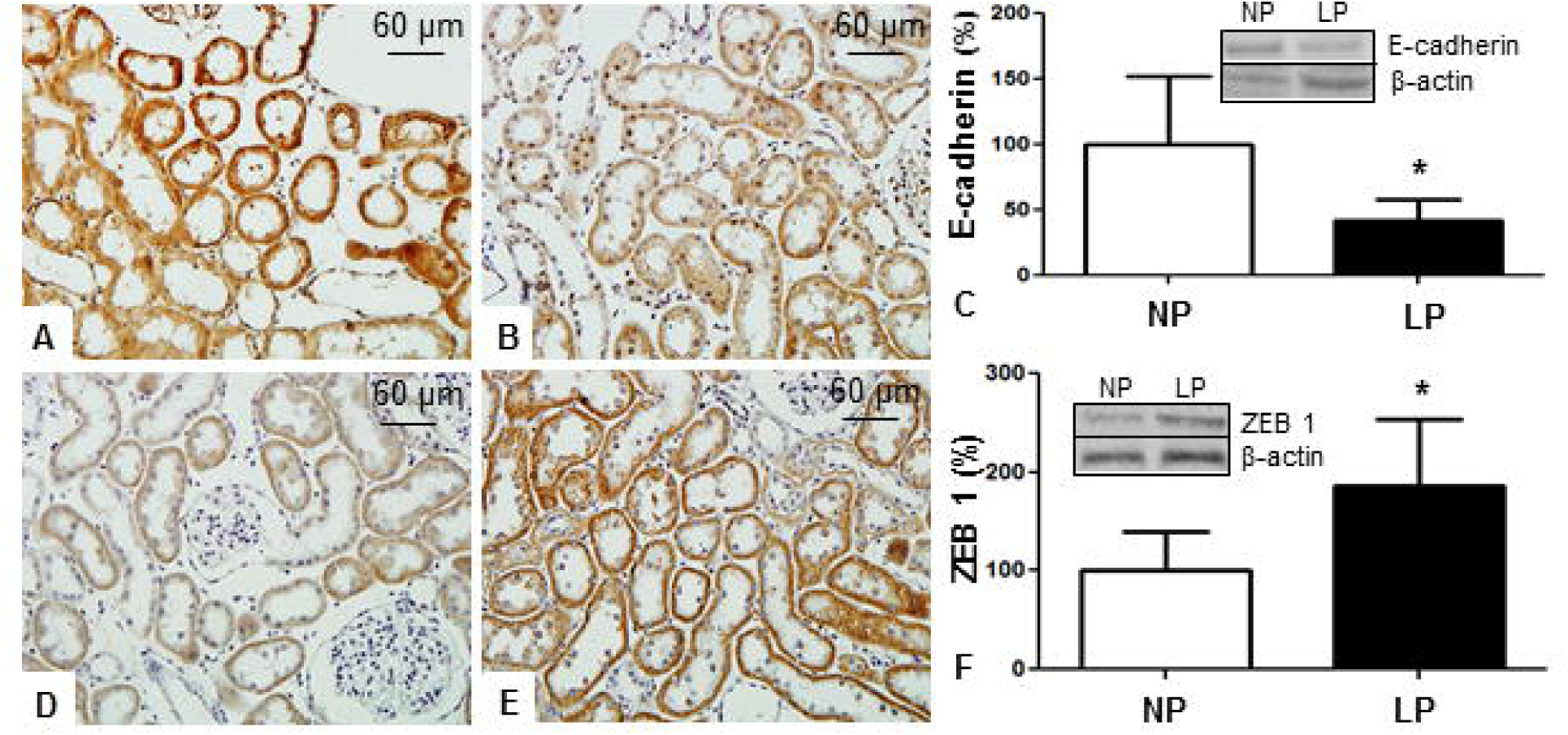
In LP (B) we found reduced E-cadherin immunoreactivity compared to that observed in NP (A) and by western blot (C) this difference was significant. Conversely, ZEB1 expression was enhanced in LP (D) in relation to NP (E). By western blot we verified that both differences were significant.

**Figure 7.**
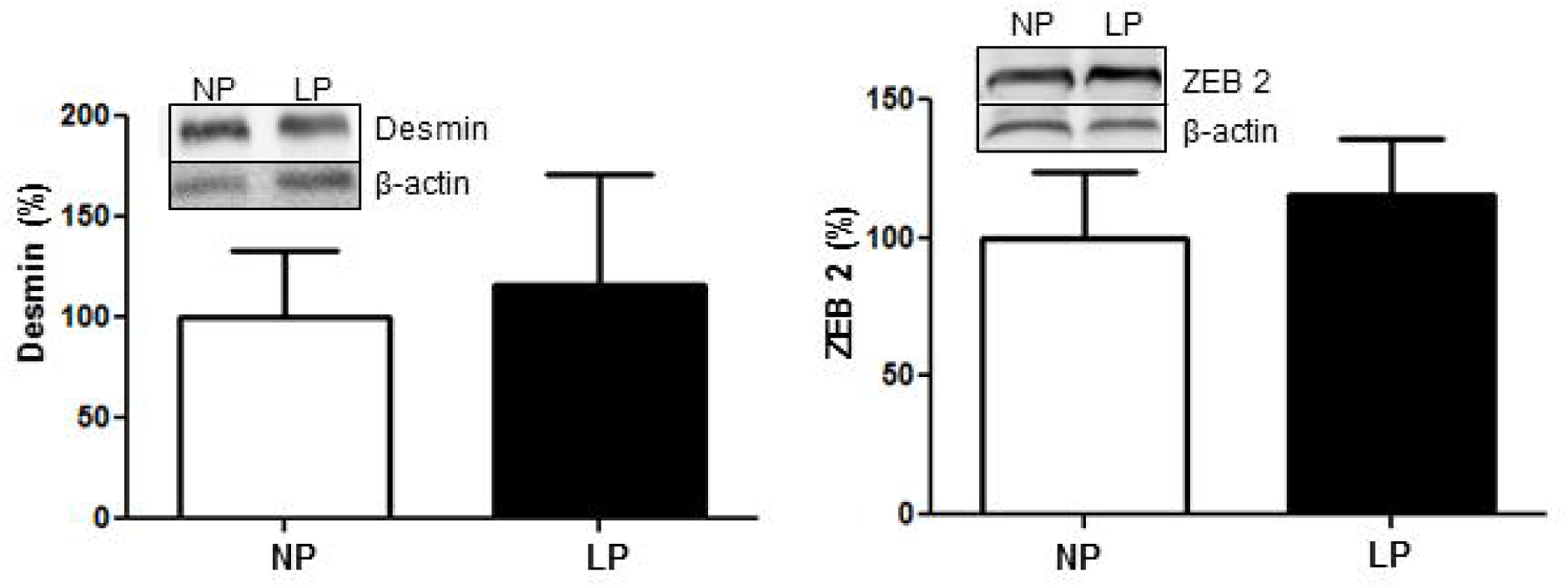
Desmin and ZEB2 western blot expression.

## DISCUSSION

More evidence is emerging that highlights the far-reaching consequences of maternal low-protein intake on kidney morphology and functional disorders. In the current study, confirming the prior reports [4,5], we demonstrated a reduction of 16% and 12% of LP offspring body weight, at birth and after 12 days of life, respectively. At this time, the kidneys were also lighter (about 14%) in LP when compared to age-matched NP offspring. However here, no difference in body and kidney mass was observed in both groups at 16 week of age. Previously, we have showed, using the same model, a reduced nephron number at 12 days (28%) and 16-wk of age (27%) accompanied by enlarged remaining glomeruli in male LP offspring [4]. At the same study, we also present a significant rise in the arterial blood pressure beyond 12-wk of age in LP offspring associated with changes in renal nerve activity and in the tubular sodium handling that lead to decreased urinary sodium excretion [4,5,7]. A few years ago Brenner *et al*. (1998) proposed that congenital reduction in nephron number become some individuals susceptible to enhanced blood pressure and kidney injury [22].

As mentioned above, the nephron-underdosing hypothesis postulated that the reduced number of nephrons contributes to arterial hypertension [4,5,22]. In this way, the current study demonstrated that, even when associated with decreased nephron number units, maternal food-restricted offspring keep a whole kidney with normal glomerular filtration rate (GFR) estimated by creatinine clearance, suggesting, in fact, a compensatory glomerular hyperfiltration despite a loss of efficiency on the filter barrier. The present investigation also demonstrated a pronounced decrease in fractional urinary sodium excretion in LP rats. The decreased FENa+ occurred in association with unchanged CCr and in the sodium-filtered load. The precise mechanism of these phenomena remains unknown. However, these results sustain the hypothesis raised urinary sodium retention and high blood pressure following maternal protein restriction, could be associated with overflow and hyperfiltration of the remaining nephrons, which in turn, could be the preponderant trigger for the development of altered glomerular ultrastructure, proteinuria and interstitial fibrosis process in LP offspring. Thereby, in the current study, the renal dysfunction and the increased blood pressure in maternal low protein diet-intake model, corroborates to confirm Brenner’s hypothesis, by which, the hyperfiltration in gestational protein-restricted offspring associated with low birthweight (LBW), leads to glomerular hypertension, proteinuria and, in the future, to sustained renal function disorder and, consequently fetal programming [4,5,20,22].

However, a great deal remains to be learned about nephron disorder in primary renal impairment of maternal protein-restricted offspring. It seems likely that glomerular and tubular structure change together, as originally suggested by Bricker [23] and Gottschalk [24]. Although in 16-wk-old LP rats we may not exclude the possibility that proteinuric glomerular disorder developed in response to an antecedent tubular or interstitial injury, nowadays, reports have shown strong evidences that once established, proteinuric glomerular injury may cause tubular dysfunction, particularly, in the current experimental model [4,5,20]. Podocytes are terminally differentiated cells incapable of regenerative postnatal replication, with major and foot process interlinked by ultrathin slit diaphragms. Podocyte injury underlies most forms of proteinuric kidney diseases [25] and, is an essential feature of progressive kidney diseases [26]. Therefore, the loss of podocytes may lead to GBM “bare” areas, which represent potential starting-point for irreversible glomerular injury [6,20,27,28]. Our group has previously showed a striking podocyte structural alteration in parallel with proteinuria and, enhanced glomerular desmin expression, denoting a decreased efficiency on the filtration barrier in 16-wk-old LP offspring when compared with age-matched control offspring. However, the mechanism by which increased filtrated protein from diseased glomeruli into the tubular lumen causes cell injury has not been entirely clear.

Here, in a gestational low-protein-treated model, we also focus on an epithelial-to-mesenchymal transdifferentiation process as a novel mechanism that promotes renal fibrosis. In the present study we investigated whether known causes of renal fibrosis TGFβ-1 act through this pathway. By immunohistochemistry, the present study verified, in 16-wk-old LP offspring, a striking enhanced entire kidney (cortical and medullar) expression of TGF-β1, fibronectin and type I collagen, intrinsically related to the fibrotic process. The current study also shows that nearly of LP offspring has increased expression (about 90%) of these proteins in the whole kidney tissue. We may state that, at least in part, the kidney expression of fibrotic and EMT markers are associated with later adult renal function disorder as an outcome, suggesting that the kidney is an organ in which fetal programming by maternal LP intake may underlie early loss of organ function and occurrence of chronic kidney disease. Simultaneously, the study shows an increase in the expression of ZEB1 in LP whole kidney by immunoblotting and immunohistochemistry at 16-week of age accompanied by a fall in the E-cadherin expression and followed by increased collagen deposition. We speculate whether the signaling receptors for specific bioactive proteins (such as TGF-β) on the tubular cell surface segments might be activated, when increased amounts of those proteins are filtered or produced by damaged nephron cells. In this way, *in vitro* experiments have shown that activation of receptors for TGF-β1 stimulates tubular cell production of inflammatory mediators [29,30,31]. In culture of immortalized rodent kidney cells, Li et al. (2008) showed that after TGF-β1 exposition, there was also loss of epithelial markers, such as E-cadherin, and acquisition of mesenchymal markers, such as collagen I and fibronectin [11].

In the current study, gene expression study confirms the enhanced renal collagen 1α2 mRNA expression, which was accompanied by increased whole kidney mRNA expression of TGF-β1, ZEB1 and fibronectin in parallel to decreased expression of E-cadherin and no change of desmin mRNA expression in 16-wk old LP offspring compared to age-matched NP rats. These results indicate that renal glomerular and tubular cells, undergo phenotypic conversion, characterized by a loss of epithelial-specific markers and a gain of transitional features, a process reminiscent of EMT [20]. As show in the current study as well as in the previous one [20], LP offspring kidneys showed enhanced expression of matrix markers (close to 80%). Results from studies in many kinds of cellular systems have been demonstrated the implication of ZEB1 as a common EMT upregulated phenomena triggered by overexpression of TGF-β1 [32-37]. Interestingly here, ZEB1 was overexpressed in 16-wk-old LP offspring entire kidney, except in glomeruli, suggesting its implication as a critical factor in the induction and/or maintenance of tubulointerstitial EMT. Thus, we may suppose that TGF-β1 triggers the tubular EMT and, its expression is up regulated in virtually every type of chronic kidney disease, including LP programming model [20,38,39].

The miRNAs are small non-coding RNA of about 21 nucleotides that regulate gene expression in a posttranscriptional manner. Our and prior study suggested that TGF-β1 pathway promotes renal fibrosis by inducing renal miRNA expression [11,20]. As miRNAs have been suggested as playing a key role in a variety of kidney diseases, we investigated whether changes in the expression of the miR-200 family and miR-192 in entire kidney from 12-days and 16-wk old LP compared to age-matched NP offspring, might also be involved in the renal pathogenesis of developmental abnormalities. At 12-days of life, we have not altered expression of these miRNAs suggesting that miRNAs expression are not programmed during pregnancy and/or are not expressed early, close to delivery. Conversely, this study has shown that in 16-wk old LP offspring, the renal tissue miR-141 (160%), miR-200c (160%) and miR-429 (163%) were significantly upregulated when compared to that observed in age-matched NP. In parallel, the mRNA for col1 α1/2 is also significantly increased (215 and 153%, respectively). These miRNAs changes, in 16-wk-old LP, occurs in parallel to the enhanced expression of ZEB1 and TGF-β1, known inducer of EMT in epithelial cells [11,30,31], and associated with unchanged ZEB2 expression, an EMT-inducing transcriptional factor essential to maintain the normal epithelial phenotype [12,13,35,37]. By immunohistochemistry, the study showed a raised type 1 collagen immunoreactivity, more pronounced in glomeruli and in corticomedullary zone of kidneys. TGF-β1 protein expression is also enhanced in the similar pattern that observed to type 1 collagen but the TGF-β1 mRNA is not altered.

Surprisingly, the present data do not supported by findings showing that renal glomeruli and tubule injury associated with fibrosis have been related to downregulation of specific miRNAs [20,40-43]. Studies have demonstrated that miR-200 family members were clearly downregulated in cells that had undergone EMT in response to TGF-β, and the expression of the miR-200 family alone was sufficient to prevent TGF-β-induced EMT [12,13]. Also, in contrast to the current findings, previous study from our laboratory and Xiong and colleagues reports verified a downregulation of the miR-200 family induced by TGF-β1 respectively, in isolated glomeruli preparation and kidney cell culture [20,31]. Thus, although members of miR-200 family have been implicated in inhibition of EMT in tubular cells, partially mediating through E-cadherin restoration at initiation of EMT [12,13] in the present study, we demonstrated a significant enhancement in the expression of mesenchymal protein markers, including fibronectin, collagen 1α1 and collagen 1α2. However, the apparent controversial results of the current study, are sustained by prior reports that have shown an overexpression of miR-192 associated with renal fibrosis induced by TGF-β1 [29,32,43,44]. Also Wang et al (2010) data demonstrated that intrarenal expression of miR 200a/b, 141, 429, 205 and 192 were increased in hypertensive nephrosclerosis and that degree of upregulation is correlated with renal disease severity [14,44]. Li et al., (2015) demonstrate the miRNAs profile of renal fibrosis and EMT, in proximal tubular cell submitted to TGF-β1 treatment, in particular by inducing renal miR-21 and miR-433 expression [43,44]. It has also been reported that there is differential transcription of miR-205 and miR-192 in IgA nephropathy, and these changes correlate with disease severity and progression [44]. On the other hand, Krupa et al., (2010) have been showed that, in proximal tubular cells stimulated by TGF-β1, expression of miR-192 decreased. Though it is clear that elevated levels of miR-192 result in a pathogenic state, the data from Krupa et al. suggest that miRNA may have a dual role and that an increase in miR-192 levels may actually be protective under certain conditions [45]. Interestingly, these authors found that a number of microRNA (miR-633, miR-34a, miR-132, miR-155) were upregulated conversely, others (miR-15a, miR-20b, miR-29c, miR-1303, miR-143 and miR-129-5p) were down-regulated.

These controversial findings highlight to complex nature of miRNA research, particularly in fetal programming models, in which many doubts persist. Notably, in previous study using the same experimental model, we have shown increased collagen deposition in isolated glomeruli, despite unchanged ZEB1 expression and conversely, glomerular ZEB2 overexpression in16-wk-old LP relative to NP offspring [20]. These results let us to speculate that ZEB1/2 expression may have specific function in different kidney structures. Comparing with prior study [20], we showed that miRNA expression pattern was widely diverse in isolated glomeruli when compared to whole kidney expression (see Figure 8). Otherwise, the gene and protein data obtained in entire kidney studies do not, necessarily, reflect the expression presented in isolated nephron segments, in fact, this observation become mandatory the restricted and isolated analyze of glomeruli and tubule structures. Taking in account that glomeruli, isolated nephron segments and interstitium are composed by a variety of cellular types, we may suppose that miRNAs, mRNAs and proteins expression profile could be deeply distinct in each nephron region. On this way, Kato et al. (2009) have considered the hypothesis that effects of renal miRs may be cell type-specific, and miR signaling networks mediates the effects of TGF β on different EMT cell types may be not the same [46]. In the present study, we could not exclude the possibility of a post-transcriptional phenomenon in the gene pathway to reduced E-cadherin protein expression. Taking in account the above results, at least in part, we may hypothesize that elevated expression of tubulointerstitial matrix markers in programmed rats, indicate that kidney cells have adopted a mesenchymal phenotype, with profound change in their morphology and function.

**Figure 8.**
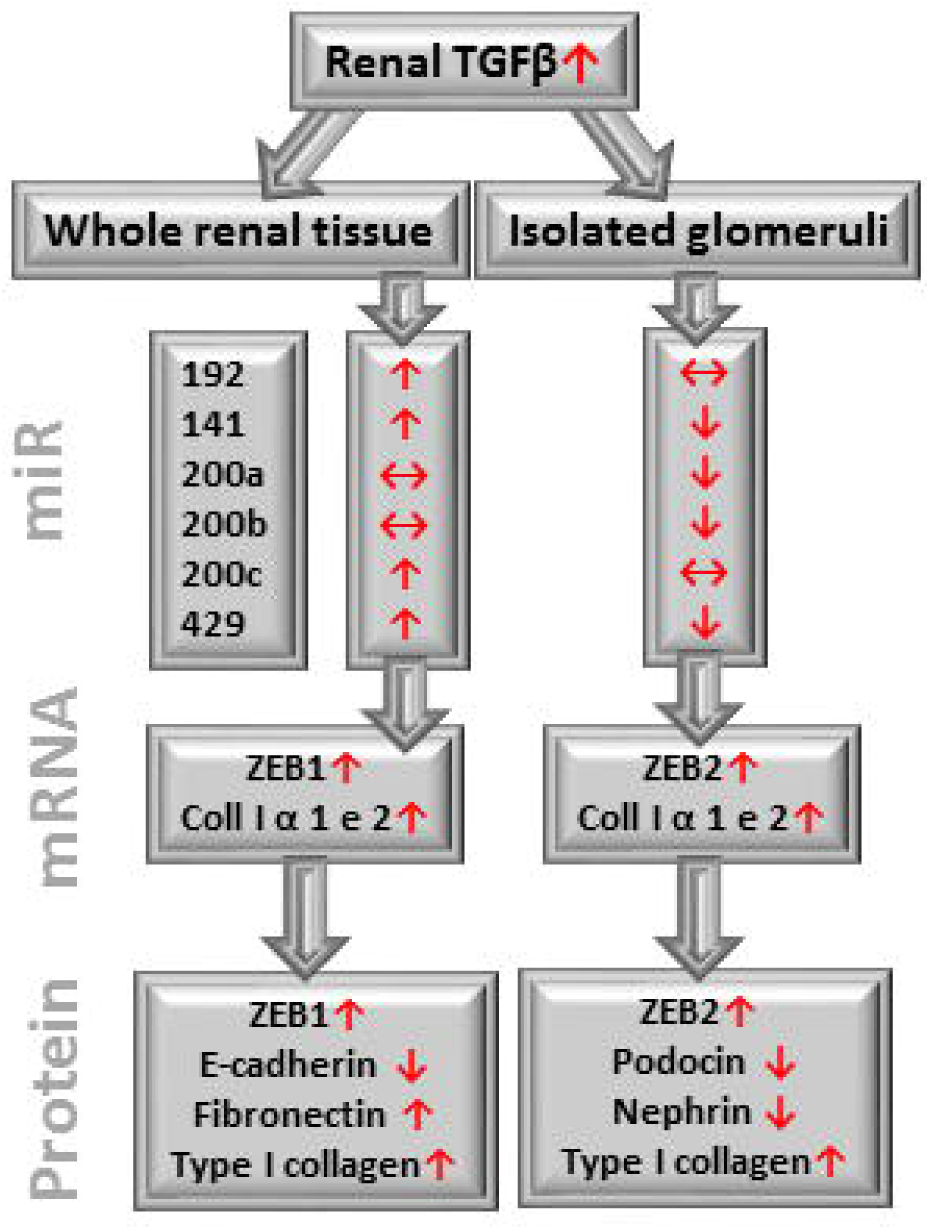
Schematic representation of fetal programming consequences in the isolated renal corpuscles compared to whole kidney in 16-week-old LP compared to NP offspring.

In the current study, we may assume that arterial hypertension and long-term hyperfiltration process manifests, itself by proteinuria was accompanied by increased whole kidney mRNA expression of TGF-β1, ZEB1, type I collagen and, fibronectin in parallel to decreased expression of E-cadherin in 16-wk old LP offspring. Surprisingly, the renal tissue miR-129, miR-141, miR-200c and miR-429 were significantly upregulated in LP offspring compared to age-matched NP rats. Despite advances, our study was not completely able to establish the precise role of miRNAs on programmed LP kidney disorders, remaining this challenger in this fertile research area, to be explored [47]. Thus, further studies are required to assess the contribution of miRNAs on renal injury progression in gestational protein-restricted model of fetal programming.

## Acknowledgments

This work was supported by Fundação de Amparo à Pesquisa do Estado de São Paulo (#09/54141-9 and #13/12486-5), CNPq and Coordenação de Aperfeiçoamento de Pessoal de Nível Superior (CAPES).

## References

1. Langley-Evans SC. Developmental programming of health and disease. Proc Nutr Soc. 2006; 65(1):97–105.

2. Langley-Evans SC. Nutritional programming of disease: unravelling the mechanism. J Anat. 2009; 215(1):36–51.

3. Barker D. The midwife, the coincidence, and the hypothesis. BMJ 2003; 20; 327(7429): 1428–30.

4. Mesquita FF, Gontijo JA, Boer PA. Expression of renin-angiotensin system signaling compounds in maternal protein-restricted rats: effect on renal sodium excretion and blood pressure. Nephrol Dial Transplant 2010a; 25: 380–388.

5. Mesquita FF, Gontijo JA, Boer PA. Maternal undernutrition and the offspring kidney: from fetal to adult life. Braz J Med Biol Res 2010b; 43: 1010–1018.

6. Rizzi VH, Sene LD, Fernandez CD, Gontijo JA, Boer PA. Impact of long-term high-fat diet intake gestational protein-restricted offspring on kidney morphology and function. J Dev Orig Health Dis. 2017; 8(1):89–100.

7. Custódio AH, de Lima MC, Vaccari B, Boer PA, Gontijo JAR. Renal sodium handling and blood pressure changes in gestational protein-restricted offspring: Role of renal nerves and ganglia neurokinin expression. PLoS One. 2017; 12(6):e0179499.

8. Kim JH, Kim BK, Moon KC, Hong HK, Lee HS. Activation of the TGF-ß/Smad signaling pathway in focal segmental glomerulosclerosis. Kidney Int 2003; 64: 1715–1721.

9. Bartel DP. MicroRNAs: target recognition and regulatory functions. Cell. 2009; 136(2):215–33.

10. Shivdasani RA. MicroRNAs: regulators of gene expression and cell differentiation. Blood. 2006; 108(12): 3646–3653.

11. Li Y, Kang YS, Dai C (2008) Epithelial-to-mesenchymal transition is a potential pathway leading to podocyte dysfunction and proteinuria. Am J Pathol 2008; 172: 299–308.

12. Gregory PA, Bert AG, Paterson EL, Barry SC, Tsykin A, Farshid G, Vadas MA, Khew-Goodall Y, Goodall GJ. The mir-200 family and mir-205 regulate epithelial to mesenchymal transition by targeting zeb1 and sip1. Nat. Cell. Biol. 2008; 10:593–601.

13. Gregory PA, Bracken CP, Bert AG, Goodall GJ. MicroRNAs as regulators of epithelial-mesenchymal transition. Cell Cycle 2008; 7: 3112–3118.

14. Wang B, Herman-Edelstein M, Koh P, Burns W, Jandeleit-Dahm K, et al. E-Cadherin Expression Is Regulated by miR-192/215 by a Mechanism That Is Independent of the Profibrotic Effects of Transforming Growth Factor-β. Diabetes 2010; 59: 1794–1802.

15. Kalluri R and Weinberg RA: The basics of epithelial mesenchymal transition. J Clin Invest 2009; 119: 1420 1428.

16. Liu Y: Epithelial to mesenchymal transition in renal fibrogenesis: Pathologic significance, molecular mechanism and therapeutic intervention. J Am Soc Nephrol 2004; 15: 1 12.

17. Thiery JP, Sleeman JP. Complex networks orchestrate epithelial-mesenchymal transitions. Nat Rev Mol Cell Biol 2006; 7(2):131–42.

18. Liu Y: Cellular and molecular mechanisms of renal fibrosis. Nat Rev Nephrol 2011; 7: 684 696.

19. Huang Y1, Tong J2, He F2, Yu X3, Fan L2, Hu J2, Tan J2, Chen Z1. miR-141 regulates TGF-β1-induced epithelial-mesenchymal transition through repression of HIPK2 expression in renal tubular epithelial cells. Int J Mol Med. 2015; 35(2): 311–318.

20. Sene L de B, Mesquita FF, de Moraes LN, Santos DC, Carvalho R, Gontijo JA, Boer PA. Involvement of renal corpuscle microRNA expression on epithelial-to-mesenchymal transition in maternal low protein diet in adult programmed rats. PLoS One. 2013; 8(8):e71310.

21. Ruijter JM, Ramakers C, Hoogaars WMH, et al. Amplification efficiency: linking baseline and bias in the analysis of quantitative PCR data. Nucleic acids research 2009; 37: e45.

22. Brenner BM, Garcia DL, Anderson S. Glomeruli and blood pressure: Less of one, more the other? Am J Hypertens 1998; 1: 335–347.

23. Bricker NS, Morrin PA, Kime SW Jr. The pathologic physiology of chronic Bright’s disease. An exposition of the “intact nephron hypothesis”. Am J Med. 1960; 28:77–98.

24. Gottschalk CW. Function of the chronically diseased kidney. The adaptive nephron. Circ Res. 1971; 28(5: Suppl 2):1–13.

25. Mundel P, Kriz W (1995) Structure and function of podocytes: an update. Anat Embryol (Berl) 1995; 192: 385–397.

26. Pippin JW, Brinkkoetter PT, Cormack-Aboud FC, Durvasula RV, Hauser PV, et al. Inducible rodent models of acquired podocyte diseases. Am J Physiol Renal Physiol 1999; 296: F213–229.

27. Fan Q, Xing Y, Ding J, Guan N, Zhang J. The relationship among nephrin, podocin, CD2AP, and alpha-actinin might not be a true ‘interaction’ in podocyte. Kidney Int 2006; 69: 1207–1215.

28. Villar-Martini VC, Carvalho JJ, Neves MF, Aguila MB, Mandarim-de-Lacerda CA. Hypertension and kidney alterations in rat offspring from low protein pregnancies. J Hypertens 2009; Suppl. 27: S47–51.

29. Chung ACK, Huang XR, Meng X, Lan HY. miR-192 Mediates TGF-β/Smad3-driven renal fibrosis. J Am Soc Nephrol 2010; 21: 1317–1325.

30. Bracken CP, Gregory PA, Kolesnikoff N, Bert AG, Wang J, et al. A double-negative feedback loop between ZEB1-SIP1 and the microRNA-200 family regulates epithelial-mesenchymal transition. Cancer Res 2008; 68: 7846–7854.

31. Xiong M, Jiang L, Zhou Y, Qiu W, Fang L, et al. The miR-200 family regulates TGF-β1-induced renal tubular epithelial to mesenchymal transition through Smad pathway by targeting ZEB1 and ZEB2 expression. Am J Physiol Renal Physiol 2012; 302: F369–379.

32. Kato M, Zhang J, Wang M, Lanting L, Yuan H, et al. MicroRNA-192 in diabetic kidney glomeruli and its function in TGF-beta-induced collagen expression via inhibition of E-box repressors. Proc Natl Acad Sci USA 2007; 104: 3432–3437.

33. Christoffersen NR, Silahtaroglu A, Orom UA, Kauppinen S, Lund AH. miR-200b mediates post-transcriptional repression of ZFHX1B. RNA 2007; 13: 1172–1178.

34. Hurteau GJ, Carlson JA, Spivack SD, Brock GJ. Overexpression of the microRNA hsa-miR-200c leads to reduced expression of transcription factor 8 and increased expression of E-cadherin. Cancer Res 2007; 67: 7972–7976.

35. Park SM, Gaur AB, Lengyel E, Peter ME. The miR-200 family determines the epithelial phenotype of cancer cells by targeting the E-cadherin repressors ZEB1 and ZEB2. Genes Dev 2008; 22: 894–907.

36. Burk U., Schubert J., Wellner U., Schmalhofer O., Vincan E., Spaderna S., Brabletz T. A reciprocal repression between zeb1 and members of the mir-200 family promotes EMT and invasion in cancer cells. EMBO Rep. 2008; 9: 582–589.

37. Korpal M, Lee ES, Hu G, Kang Y. The miR-200 family inhibits epithelialmesenchymal transition and cancer cells by targeting the E-cadherin transcriptional repressors ZEB1 and ZEB2. J Biol Chem 2008; 283: 14910–14914.

38. Yang J, Liu Y. Dissection of key events in tubular epithelial to myofibroblast transition and its implications in renal interstitial fibrosis. Am J Pathol 2001; 159: 1465–1475.

39. Böttinger EP, Bitzer M. TGF-beta signaling in renal disease. J Am Soc Nephrol 2002; 13: 2600–2610.

40. Ho J, Ng KH, Rosen S, Dostal A, Gregory RI, Kreidberg JA. Podocyte-specific loss of functional microRNAs leads to rapid glomerular and tubular injury. J Am Soc Nephrol. 2008; 19(11): 2069–2075.

41. Shi S, Yu L, Chiu C, Sun Y, Chen J, Khitrov G, Merkenschlager M, Holzman LB, Zhang W, Mundel P, Bottinger EP. Podocyte-selective deletion of dicer induces proteinuria and glomerulosclerosis. J Am Soc Nephrol 2008; 19: 2159–2169.

42. Wang Q, Wang Y, Minto AW, Wang J, Shi Q, Li X, Quigg RJ. MicroRNA-377 is up-regulated and can lead to increased fibronectin production in diabetic nephropathy. FASEB J 2008; 22: 4126–4135

43. Li Y, An H, Pang J, Huang L, Li J, Liu L. MicroRNA profiling identifies miR-129-5p as a regulator of EMT in tubular epithelial cells. Int J Clin Exp Med 2015; 8(11):20610–6.

44. Wang B, Koh P, Winbanks C, Coughlan MT, McClelland A, et al. miR-200a prevents renal fibrogenesis through repression of TGF-{beta}2 expression. Diabetes 2011; 60: 280–287.

45. Krupa A, Jenkins R, Luo DD, Lewis A, Phillips A, Fraser D. Loss of MicroRNA-192 promotes fibrogenesis in diabetic nephropathy. J. Am. Soc. Nephrol 2010; 21(3), 438–447.

46. Kato M, Arce L, Natarajan R (2009) MicroRNAs and their role in progressive kidney diseases. Clin J Am Soc Nephrol 2009; 4: 1255–1266.

47. Carew RM, Wang B, Kantharidis P. The role of EMT in renal fibrosis. Cell Tissue Res 2012; 347: 103–116.

